# Microencapsulation of mesenchymal stromal cells in covalent alginate hydrogels for cell therapy

**DOI:** 10.1101/2023.11.27.568852

**Authors:** Mathilde Ambrosino, Fabien Nativel, Cécile Boyer, Nathan Lagneau, François Loll, Boris Halgand, Farida Djouad, Denis Renard, Arnaud Tessier, Jérôme Guicheux, Vianney Delplace, Catherine Le Visage

**Author notes:** co-authors (Vianney Delplace, Catherine Le Visage).

## Abstract

Osteoarthritis (OA) is the most common inflammatory joint disease and currently lacks an effective curative treatment. Intra-articular injection of mesenchymal stromal cells (MSCs) has gained attention as a relevant therapeutic approach for OA treatment due to the MSC’s ability to secrete anti-inflammatory and immunomodulatory factors. Given their limited viability post- intraarticular injection and the potential leakage of cells out of the injection site, encapsulating MSCs in hydrogels is considered a promising strategy to protect them and provide a suitable 3D microenvironment to support their biological activities. Calcium-cross-linked alginate hydrogels are commonly used for MSC encapsulation, but their long-term in vivo stability remains uncertain. On the other hand, alginate cross-linking by the strain-promoted azide-alkyne cycloaddition (SPAAC) reaction would create a network unaffected by an ionic environment. Hence, this study aimed to develop an alginate-based hydrogel cross-linked via stable and cytocompatible covalent bonds for cell encapsulation. We established for the first time the formation of covalent alginate hydrogels between two SPAAC precursors, namely alginate-BCN and alginate-N_3_. These hydrogels exhibited in vitro stability and enabled the diffusion of molecules of interest. We then generated alginate-based SPAAC microgels of 170 μm in mean diameter, suitable for intra-articular injection. We next encapsulated human adipose MSCs (hASCs) in these alginate-based SPAAC microgels and confirmed their cytocompatibility, with over 90 % of cells remaining viable after 14 days in culture. Finally, the microencapsulated hASCs maintained their biological properties and were able to secrete anti-inflammatory factors (IDO, PGE2, and HGF) when exposed to pro-inflammatory cytokines (TNF-α and IFN-γ). In the end, human-activated lymphocytes were cultured in contact with microencapsulated hASCs, and CD3+ T cell proliferation was quantified by flow cytometry. We demonstrated that the encapsulation process did not impair the hASC immunomodulatory activity. Overall, our findings show the potential of alginate-based SPAAC hydrogels for microencapsulating hASCs for cell therapy.

## 1 Introduction

Osteoarthritis (OA) is the most widespread joint disease globally, impacting 7 % of the global population, representing more than 500 million people [1]. OA is a multifactorial condition driven by various factors, including genetics, environment, and lifestyle. It begins with a disturbance at the molecular level, marked by abnormal metabolism of joint tissues. Then, it progresses to anatomical and/or physiological disorders, resulting mainly in cartilage degradation, inflammation of the synovial membrane, and bone remodeling, leading to the establishment of the disease [2]. The burden of OA on patients is profound, causing pain, functional impairment, and a diminished quality of life. Despite its prevalence, OA treatments remain challenging, with current interventions primarily focused on symptom management [3]– [6]. Consequently, there is an urgent need for innovative therapies capable of slowing down or stopping OA progression.

Mesenchymal stromal cells (MSCs) delivery has emerged as a promising approach in cell therapy and regenerative medicine. Initially identified and isolated from bone marrow, MSCs can now be obtained from various tissues, including adipose tissue, offering practical advantages such as accessibility, differentiation potential, and proliferative capacity [7], [8], [9]. MSCs are particularly attractive for OA treatment due to their ability to secrete anti-inflammatory and immunomodulatory factors such as Indoleamine 2,3-dioxygenase (IDO), Prostaglandin E2 (PGE2), Human Growth Factor (HGF) and Transforming Growth Factor-β (TGF-β) [10], [11]. Pre-clinical and clinical studies involving intra-articular (IA) injections of MSCs have demonstrated their potential to slow down cartilage destruction, reduce inflammation [12], [13], reduce pain, and improve joint functionality [14]–[17]. However, short-term challenges like cell migration from injection sites [18], [19] and cell death due to mechanical stress [20] have hindered their therapeutic efficacy.

Cell encapsulation techniques have been explored to address these challenges, aiming to enhance the longevity and effectiveness of MSC-based OA therapies. Hydrogels, which are hydrated polymer networks, could protect cells during and after their injection, create a favorable environment for their survival, and maintain them within the joint to prolong their therapeutic effects. Alginate, a well-studied natural polymer that can be ionically cross-linked with calcium, has demonstrated potential for the encapsulation of adipose-derived MSCs (ASCs) [21]–[23]. However, concerns remain regarding the long-term stability of these ionic alginate hydrogels due to the presence of monovalent ions in the synovial liquid, which could eventually lead to progressive solubilization of this ionic network. Hydrogel stability is crucial to prolong MSCs’ presence in the joint cavity, maximize their ability to release therapeutic factors in response to OA-related inflammation, and reduce the need for multiple administrations to the patient.

Therefore, we explored an alternative approach and developed a new alginate-based system that has not been documented yet. This system involves the covalent cross-linking of alginate using the strain-promoted azide-alkyne cycloaddition (SPAAC) reaction between a bicyclononyne (BCN) and an azide (N_3_). SPAAC reactions are notable for satisfying both click and bioorthogonal chemistry criteria, ensuring no interference with biological components [24], [25]. We previously reported using SPAAC with hyaluronic acid modified with BCN or N_3_, rapidly generating covalent and versatile hydrogels. Human ASCs (hASCs) were encapsulated in these hyaluronic acid-based hydrogels, and their cytocompatibility and effects on encapsulated cell secretions were confirmed [26]. In this study, creating micro-sized hydrogels was crucial for developing a cell therapy, enabling the intra-articular injection of alginate-based SPAAC hydrogels through a needle. It is worth mentioning that, to our knowledge, alginate-based SPAAC hydrogel micromolding has not been previously attempted.

This study aimed to encapsulate hASCs in alginate cross-linked by covalent bonds via the SPAAC reaction for potential application in OA cell therapy. We modified alginate chains by introducing BCN or N_3_ groups in a one-step synthesis. Pre-hydrogel solutions were then mixed to create stable alginate-based SPAAC hydrogels. We then demonstrated the diffusion of molecules of interest through these hydrogels by a release monitoring method. Using a micromolding protocol developed in our laboratory [21], we successfully generated alginate-based SPAAC microgels with an average diameter of 170 µm. After confirming the feasibility of encapsulating hASCs in these alginate-based SPAAC microgels, we confirmed their cytocompatibility, with more than 90 % of cells remaining viable after 14 days of culture. From there, we explored the secretory activity of the microencapsulated cells when treated with pro-inflammatory factors. We demonstrated that microencapsulated hASCs maintained their ability to secrete anti-inflammatory compounds. Finally, we co-cultured microencapsulated hASCs with human activated lymphocytes and analyzed CD3+ T cell proliferation. We demonstrated that the encapsulation process did not impair the cell immunomodulatory activity.

## 2 Materials and methods

### 2.1 Materials

Sodium alginate powder (Protanal™ LF10/60FT, 60–180 kDa, 25–35 % mannuronic acid, and 65–75 % guluronic acid) was purchased from FMC Biopolymer. N-[(1R,8S,9s)- Bicyclo[6.1.0]non-4-yn-9-ylmethyloxycarbonyl]-1,8-diamino-3,6-dioxaoctane (BCN), Cell Counting Kit – 8 (CCK-8), alginate lyase powder and fluorescein sodium-salt were obtained from Sigma-Aldrich. Azido-PEG2-amine was purchased from BroadPharm. 2-(N-morpholino)ethanesulfonic acid (MES) and 4-(4,6- dimethoxy-1,3,5-triazin-2-yl)-4- methylmorpholinium chloride (DMT-MM) were purchased from TCI Chemicals. Polydimethylsiloxane (PDMS, RTV 615, used as a 2-part kit with a 10:1 mixing ratio) was obtained from Eleco Produits. Fetal calf serum (FCS) and lymphocyte separation medium were purchased from Eurobio Scientific. Mesenchymal stem cell growth medium 2 was purchased from PromoCell. Violet proliferation dye 450 was obtained from BD biosciences. Phytohemagglutinin-L and Iscove’s Modified Dulbecco’s Media were obtained from Life Technologies. µ-slide 8 wells were purchased from ibidi. PGE2 ELISA Kit-Monoclonal was obtained from Cayman Chemical, following the manufacturer’s instructions. Human TNF-α and human IFN-γ were purchased from Miltenyi Biotec. Human TNF-α DuoSet ELISA, Human IFN-γ DuoSet ELISA, human HGF DuoSet ELISA, and human TGF-β1 DuoSet ELISA were obtained from R&D Systems, following the manufacturer’s recommendations. Otherwise stated, all other reagents were purchased from ThermoFisher Scientific.

### 2.2 Methods

#### 2.2.1 Component synthesis and alginate-based SPAAC hydrogel formation

##### Synthesis of alginate-BCN

Alginate-BCN was synthesized following a previously reported protocol with minor modifications [26]. Alginate (100 mg) was dissolved in MES buffer pH 5.5 (9 mL) and stirred at room temperature. DMT-MM (28 mg, 0.4 eq) was added and allowed to react for 30 minutes. Subsequently, BCN (16 mg; 0.2 eq) was dissolved in 1 mL of DMSO, and the solution was added dropwise. The reaction was carried out under continuous stirring for 3 days at room temperature. Then, the solution was dialyzed against 0.1 M DPBS containing an additional 2 % DMSO overnight, against 0.1 M DPBS for 8 hours, and against deionized water for 2 days. After dialysis, the solution was sterilized on a 0.22 µm filter, lyophilized, and stored at -80°C.

The degree of substitution of alginate-BCN was determined by elementary analysis using the N/C ratio and the equation previously reported in [27].

##### Synthesis of alginate-N_3_

Alginate-N_3_ was synthesized as described above, with minor modifications. Alginate (100 mg) was dissolved in MES buffer pH 5.5 (9 mL) and stirred at room temperature. DMT-MM (70 mg, 1 eq) was added and allowed to react for 30 minutes. Subsequently, azido-PGE2-amine (22 mg; 0.5 eq) was dissolved in 1 mL of MES buffer pH 5.5, and the solution was added dropwise. The reaction was carried out under continuous stirring for 3 days at room temperature. Then, the solution was dialyzed against 0.1 M DPBS for 1 day and against deionized water for 2 days. After dialysis, the solution was sterilized on a 0.22 µm filter, lyophilized, and stored at -20°C. The degree of substitution of alginate-N_3_ was determined by elementary analysis using the N/C ratio and the equation previously reported in [27].

##### Hydrogel formation

Alginate-BCN and alginate-N_3_ were separately dissolved in PBS or cell culture medium (DMEM) at room temperature, at a fixed total alginate concentration of 1% (w/v). These pre-hydrogel solutions were then mixed together, with a BCN/N_3_ molar ratio of 1:1.4, to form alginate-based SPAAC hydrogels. In all experiments, except for gelation time measurements and stability evaluation, the hydrogel formation process was conducted for 1 hour at 37°C in a humid atmosphere.

#### 2.2.2 Physicochemical characterization of alginate-based SPAAC hydrogels Gelation time measurements

The pre-hydrogel solutions were prepared as described above in PBS or DMEM, and measurements were rapidly conducted upon combining a total volume of 1 mL of the two alginate components. Rheological analysis was carried out using a HAAKE™ MARS™ rheometer (ThermoFisher Scientific) equipped with a 60 mm titanium upper cone (C60/1°, Ti L geometry) and a Peltier plate for precise temperature control. Time sweep experiments were executed while maintaining a constant stress of 1 Pa and a constant frequency of 1 Hz, ensuring operation within the linear viscoelastic range. Gelation time experiments were conducted at room temperature and 37°C, using a solvent trap to prevent evaporation. Gelation time was determined as the time when the storage modulus (G’) equaled the loss modulus (G’’). The experiments were performed three times for each condition, in either PBS or DMEM and at room temperature or 37°C.

##### Stability evaluation

The stability studies were conducted either in PBS or DMEM by adapting a previously reported procedure [26]. Briefly, a pre-hydrogel solution (200 μL) was incubated at 37°C for 10, 20, 30 minutes, 1 hour and overnight. After weighing the hydrogel and adding warm DMEM (37°C), supernatants were removed at defined time points (0, 1, 2, 4, 6, and 24 hours; then 3, 7, 14, 21, 28, and 56 days). The stability was determined by calculating the ratio of hydrogel mass to its initial mass at a particular time point. Each experimental condition in DMEM was repeated three times.

##### Compression tests

Compression experiments were performed on cylindrical hydrogels of 1 mm in height and 4 mm in diameter using a MicroTester (CellScale). Pre-hydrogel solutions were prepared as described above, in PBS or DMEM, then transferred to a 37°C incubator for 1 hour in a humid atmosphere. The compression measurements were performed under 20% strain. Each hydrogel underwent a 30-second compression cycle using a 6 mm square stainless-steel plate placed upon a 58 mm long microbeam (558.8 μm diameter). Young’s modulus was computed in accordance with the manufacturer’s guidelines, as previously reported [21], [26]. For each condition, measurements were performed on at least 4 hydrogels for repeatability assessment.

#### 2.2.3 Diffusion properties of alginate-based SPAAC hydrogels

To investigate the diffusion properties of alginate-based SPAAC hydrogels, the release of Fluorescein, FITC-conjugated BSA, TNF-α, and IFN-γ from the hydrogels was evaluated. Each molecule was dissolved in PBS at 10 µg/mL, 20 µg/mL, 200 ng/mL, and 200 ng/mL, respectively. These solutions were used to dissolve the alginate-BCN and alginate-N3 hydrogel precursors separately, as described above. Then, 200 μL of the pre-hydrogel solution, in a BCN/N_3_ molar ratio of 1:1.4, was transferred in 2 mL-Eppendorf tubes before incubation at 37°C for 1 hour for cross-linking. Warm PBS (37°C, 800 μL) was then added to investigate the release of the molecules of interest. At specific time points, supernatants were removed and stored at 4°C, and then 800 µL of warm PBS were added to maintain near-sink conditions. After the final time point, the hydrogels were dissolved by adding alginate lyase (10 U/mL) to ensure the release of any remaining molecules. The concentrations of released fluorescent molecules and cytokines in the supernatants were quantified using either a microplate reader for fluorescent measurements (Tristar 2S, Berthold Technologies) or ELISA kits (R&D Systems) following the manufacturer’s instructions. Free molecule solutions were incubated, sampled, and stored in conditions similar to the released molecules to serve as degradation controls. The relative release fraction at each time point was determined by normalizing the released molecule concentration at each time point to the free molecule control. Results are presented as the cumulative release of each molecule. Ionic alginate cross-linked with calcium (2 % w/v, 100 mM CaCl_2_) was used as a control. Each experimental condition was replicated four times.

#### 2.2.4 Micro-encapsulation of hASCs in alginate-based SPAAC microgels

##### Micromolding of alginate-based SPAAC hydrogels

Hydrogel micromolding was executed using polydimethylsiloxane (PDMS) micromolds obtained via soft lithography, as previously described [21]. PDMS micromolds were sterilized in an autoclave and hydrophilized using O_2_ plasma treatment (Zepto plasma system, Diener electronic; 40Watts, 90 s) immediately before use. First, alginate-BCN and alginate-N_3_ were dissolved in complete cell culture medium (DMEM, 10% v/v fetal calf serum (FCS), 1% v/v penicillin-streptomycin, 1% v/v fungizone), at a fixed total alginate concentration of 1% (w/v). PDMS micromolds were filled by evenly distributing the pre-hydrogel solution on the micromold surface. The volume of solution added was 200 µL for 1,600 microgels and 800 µL for 10,000 microgels. Any surplus was manually removed using a sterile glass slide. PDMS micromolds filled with the pre-hydrogel solution were then placed in a Petri dish and incubated for 1 hour at 37°C in a humid atmosphere for gelation. After cross-linking, the microgels were removed by gently scraping the micromold surface in a complete cell culture medium. The microgels were subsequently transferred to 1.5-mL Eppendorf tubes containing complete cell culture medium and stored at 37°C.

##### Microgels stability evaluation

The size and shape of the microgels were observed under light microscopy over time (immediately after microencapsulation, then 1, 3, 7, 14, and 21 days) using a Digital Microscope VHX-7000 (Keyence). The microgels were immersed in complete cell culture medium and stored at 37°C. The mean diameter of the microgels was determined using the software associated with the Digital Microscope VHX-7000. The experiments were repeated three times, with 100 microgels analyzed for each time point.

##### Human adipose-derived mesenchymal stromal cells isolation and cell culture

Human adipose-derived mesenchymal stromal cells (hASCs) were collected from the subcutaneous adipose tissue of three different donors undergoing liposuction with their informed consent, as previously described [28]. After isolation, hASCs were expanded in MSC growth medium, supplemented with 1% v/v penicillin-streptomycin and 1% v/v fungizone. For all 2D and 3D experiments, hASCs were cultured in complete cell culture medium, from passage 2 to passage 5. Cytocompatibility experiments were performed with cells from donor A, while bioactivity experiments were performed with cells from three separate donors (donors A-C, Supp. Table 1).

##### Encapsulation of hASCs in alginate-based SPAAC microgels

Sterile alginate-BCN and alginate-N_3_ were dissolved in complete cell culture medium at a fixed total alginate concentration of 1% (w/v) and a BCN/N_3_ molar ratio of 1:1.4. Cells were suspended in the pre-hydrogel solution at a seeding density of 3 million cells per mL. The cell-loaded pre-hydrogel solution was subsequently evenly dispersed over the hydrophilized PDMS micromolds. A centrifugation step (300g, 2 minutes) was conducted at room temperature to incorporate the cells. Then, cell-loaded microgels were obtained following the aforementioned protocol for alginate-based SPAAC microgels preparation. After transferring the cell-loaded microgels to 1.5 mL-Eppendorf tubes containing complete cell culture medium at 37°C, half of the culture medium was renewed every 2-3 days.

##### Cell-loaded microgels stability evaluation

The size and shape of the microgels were observed under light microscopy over time (immediately after microencapsulation, then 1, 3, 7, and 14 days) using confocal microscopy Nikon A1RSi (Nikon Champigny sur Marne). The microgels were immersed in complete cell culture medium and stored at 37°C. The mean diameter of the microgels was determined using the ImageJ software.

#### 2.2.5 Cytocompatibility of alginate-based SPAAC microgels on hASCs

##### Viability, metabolic activity, and DNA quantification

Cell viability, metabolic activity, and DNA quantification evaluations of microencapsulated hASCs were conducted on day 0, 1, 3, 7, and 14, using LIVE/DEAD™ staining, Cell Counting Kit-8 (CCK-8) assay, and PicoGreen™ assay, respectively. Cytocompatibility experiments were performed with cells from donor A (Supp. Table 1). For LIVE/DEAD™ staining, microgels containing cells were incubated with calcein-AM and ethidium homodimer-1, according to the manufacturer’s protocol. In a separate experiment, Alexa Fluor™ 568 Phalloidin and TO-PRO™-3 Iodide (642/661) staining was performed to analyze cell distribution within the microgels, following the manufacturer’s recommendations. After incubation, the microgels were transferred from the 1.5 mL-Eppendorfs into an 8-well chamber slide (ibidi) for image acquisition with confocal microscopy (A1RS, Nikon). The ImageJ software was used to calculate cell viability. In a separate experiment, the metabolic activity was evaluated by CCK-8 assay, following the manufacturer’s instructions. Microencapsulated cells were incubated for 4 hours with the CCK-8 solution, and the absorbance of the supernatant at 460 nm was measured using a microplate reader (Tristar 2S, Berthold Technologies). Then, the number of encapsulated cells was evaluated by DNA quantification using the PicoGreen™ assay, according to the manufacturer’s recommendations. To degrade alginate-based SPAAC formation and release the cells, a solution of alginate lyase (10 U/mL) was added to each Eppendorf for 15 min under agitation. After centrifugation (300 g, 5 minutes), the supernatants were removed, and cell pellets were rinsed twice with PBS before adding TE 1X buffer and subsequently freezing at -80°C until analysis. After thawing, the samples were transferred into a 96-well plate, and the PicoGreen reagent was added. DNA quantification was obtained by fluorescent measurement (ex 480 nm/em 520 nm; Tristar 2S, Berthold Technologies).

#### 2.2.6 Biological activity of microencapsulated hASCs Secretory activity

The secretory functions of microencapsulated hASCs were investigated as previously described, with minor modifications [21]. Microencapsulated cells (20,000 cells) were gradually deprived of FCS, starting at 10% on the day of encapsulation, decreasing to 5% on day 3, and further lowering to 2.5% on day 4. Cells cultured in 2D were used as controls, with 20,000 cells per well seeded in a 24-well plate. On day 4, cells were stimulated with a cell culture medium containing 1% FCS, supplemented with 20 ng/mL of TNF-α and 20 ng/mL of IFN-γ. Non-treated cells were cultured with 1% FCS, with no supplementation. After 72 hours of stimulation, supernatants were collected to assess the release of soluble factors. The concentrations of PGE2, HGF, and TGF-β in supernatants were determined using ELISA kits, following the manufacturer’s instructions. The enzymatic activity of IDO in supernatants was measured by monitoring the conversion of tryptophan to kynurenine, with a photometric determination of kynurenine concentration in the supernatants, as described before [29]. Each experimental condition was carried out in triplicate, using cells from three separate donors (donors A-C, Supp. Table 1). The experiments were repeated twice for each donor. Results were expressed as the concentration of each molecule normalized to the number of cells after 72 hours of stimulation, which was determined using the PicoGreen™ assay.

##### Immunosuppression assay

To assess the immunosuppressive properties of hASCs, peripheral blood mononuclear cells (PBMCs) were cultured alone or in contact with hASCs, control in 2D or microencapsulated.

Fresh PBMCs from healthy donors were obtained after a density gradient centrifugation using lymphocyte separation medium (1.077 g/ml). Once purified, PBMCs were labeled with violet proliferation dye 450, according to the manufacturer’s instructions, and plated in the presence or absence of hASCs at a cell ratio of 1 hASC per 10 PBMC. Cells were cultured in mixed lymphocyte reaction (MLR) medium, containing 10% inactivate FCS, 1% P/S, 1% sodium pyruvate, 1% non-essential amino acids, 1% glutamine, and 100 µM HEPES in Iscove’s Modified Dulbecco’s Media (IMDM), at 37°C in a humidified atmosphere containing 5% CO2. Lymphocyte proliferation was activated with Phytohemagglutinin-L (PHA) (5 mg/mL). After 4 days, CD3+ T cell proliferation was quantified by measuring the corresponding decrease in cell fluorescence by flow cytometry (BD FACS Symphony A5.2). The experiments were performed with cells from three separate donors (donors A-C, Supp. Table 1).

#### 2.2.7 Statistical analysis

All data are presented as mean ± standard deviation (SD). Statistical analyses were performed using GraphPad Prism 8. Statistical significance was set at p < 0.05. Compression tests and stability of microgels in DMEM were compared using one-way ANOVA tests, followed by ad-hoc tests. Unpaired t-tests were performed to compare in vitro data of PGE2, HGF, and TGF-β concentrations and IDO activity. The Kruskal-Wallis test was performed to compare in vitro data of MSCs immunomodulation on lymphocytes.

## 3 Results

### 3.1 Alginate-based SPAAC hydrogel synthesis

The SPAAC reaction, described previously in the literature [26], [30], [31], is a click and bioorthogonal reaction that has been chosen to produce covalently cross-linked alginate hydrogels. In this study, two single-step syntheses were necessary to obtain the two alginate-based SPAAC hydrogel precursors of interest, namely alginate-BCN and alginate-N_3_. The two hydrogel precursors were obtained through a standard amidation procedure utilizing DMT-MM as an activating agent under mild aqueous conditions of reaction and purification. The successful functionalization of alginate-BCN and alginate-N_3_ was confirmed by elemental analysis, revealing degrees of substitution of 7 ± 3 % and 30 ± 2 %, respectively.

### 3.2 Alginate-based SPAAC hydrogel formation and characterization

To form hydrogels, alginate-BCN and alginate-N_3_ were first dissolved separately in either PBS or DMEM at room temperature at a fixed alginate concentration of 1 % (w/v). Alginate-based SPAAC hydrogels were then obtained by mixing these two pre-hydrogel solutions in a BCN/N_3_ molar ratio of 1:1.4 based on preliminary data (Fig. 1a). We first characterized the gelation time of this alginate-based SPAAC hydrogel formulation via rheological measurements (i.e., time sweep assay) at room temperature or 37°C. A shorter gelation time, defined as the time when G’ and G’’ intersect, was evidenced when the alginate-based SPAAC hydrogel was formed at 37°C (≈ 20 minutes) compared to room temperature (≈ 50 minutes) (Fig. 1b). The much slower gelation at room temperature is advantageous because it allows the user to prepare a well-mixed polymer solution before gelation. In comparison, a faster gelation time at 37°C enables efficient cross-linking under incubation conditions for in vitro experiments. Then, we evaluated the stability of the alginate-based SPAAC hydrogels as a function of cross-linking time. After mixing the two hydrogel precursor solutions, the pre-hydrogel solution was immersed in DMEM at 37°C after a defined cross-linking time. The stability was assessed for five different cross-linking times (i.e., 10, 20, 30 minutes, 1 hour and overnight). A minimum of 30 minutes of cross-linking at 37°C was necessary to prevent hydrogel dissolution (Fig. 1c). We also observed that hydrogels cross-linked for 30 minutes were not stable over time, in contrast to hydrogels cross-linked for 1 hour or overnight at 37°C, which remained stable over 2 months (Fig. 1d). Thus, for cell encapsulation, we chose to continue our experiments with a 1-hour cross-linking time at 37°C before incubation. Finally, to evaluate the mechanical properties of the SPAAC hydrogels over time, we performed compression tests on hydrogels dissolved in

**Figure 1:**
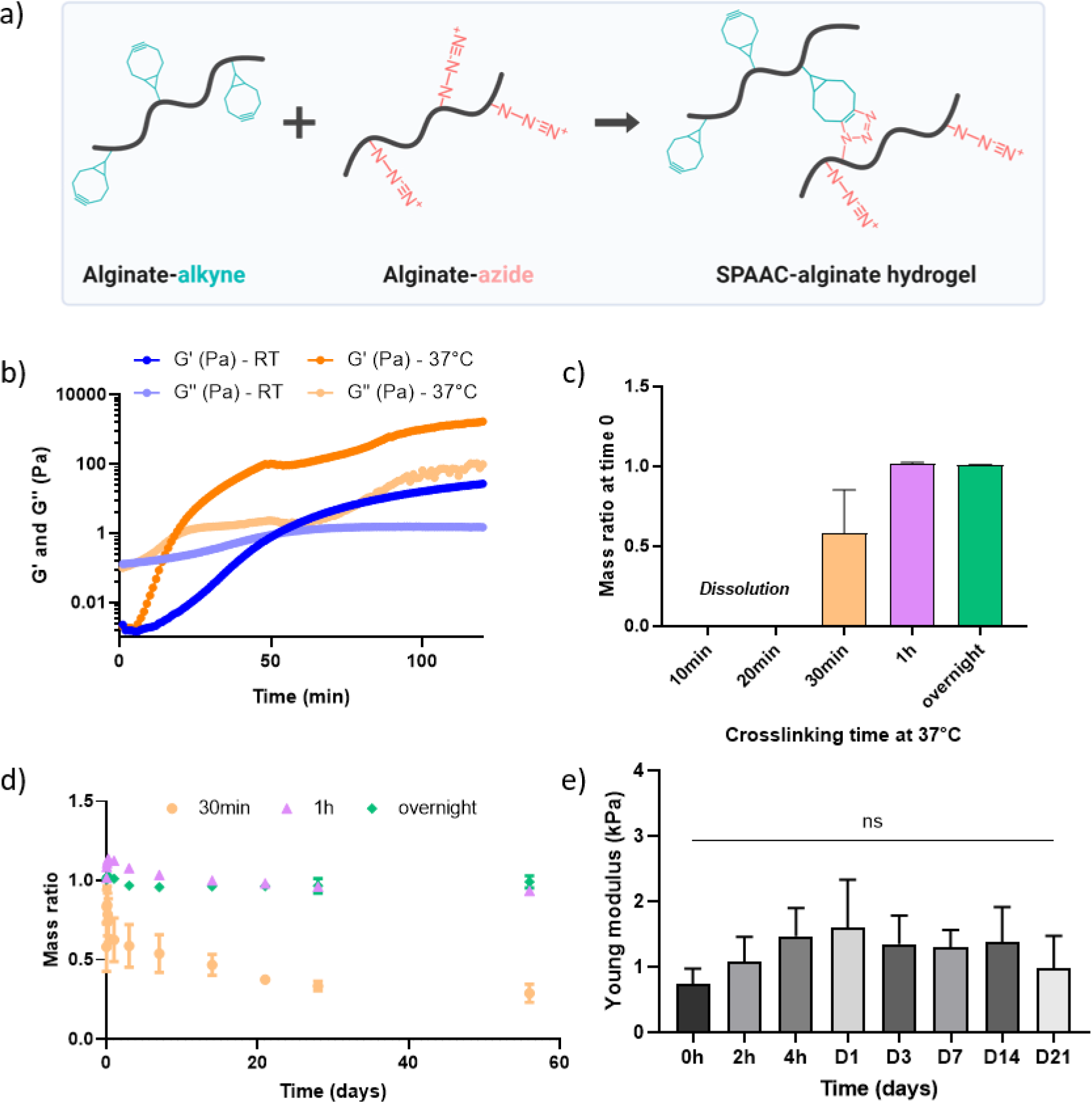
Physicochemical characterization of alginate-based SPAAC hydrogels. a) Chemical representation of the formation of alginate-based SPAAC hydrogel by combination of alginate-alkyne (BCN) and alginate-azide (N_3_); b) Time sweep experiments representing the evolution of G’ and G’’ as a function of time and temperature in DMEM (N=3, n=1); The influence of cross-linking time on hydrogel mass ratio on stability : c) in the short term, i.e., at time 0 (N=1, n=3) and d) over 2 months (N=1, n=3) in DMEM; e) Hydrogel stiffness over 21 days in DMEM, after 1 hour of cross-linking in 37°C (N=1, n=4 à 6).

DMEM and cross-linked for 1 hour at 37°C. Our results show that our alginate-based SPAAC hydrogel has a Young’s modulus of ≈ 1-2 kPa that is constant over at least 21 days. Similar studies were performed using PBS, leading to similar results (Suppl. Fig. S1).

### 3.3 Diffusion properties of alginate-based SPAAC hydrogels

The diffusion properties of the alginate-based SPAAC hydrogel were investigated, with ionic alginate as a control. In a pro-inflammatory microenvironment, pro-inflammatory cytokines (e.g., TNF-α and IFN-γ) would have to diffuse inside the hydrogel to activate the encapsulated MSCs (Fig 2a). In response to this stimulus, the encapsulated MSCs would secrete immunomodulatory molecules that would have to exit the hydrogel to exert their effects on surrounding cells. Thus, molecular diffusion is vital to a successful hydrogel-assisted immunomodulatory cell therapy. The secretome of MSCs contains a variety of molecules with different molecular weights (MW). To perform preliminary experiments on the diffusion properties of alginate-based SPAAC hydrogels, we selected two fluorescent model molecules covering a large range of MWs: a low-MW molecule (fluorescein, 370 Da) and a high-MW protein (fluorescent BSA, 66 kDa). Diffusion capacity was assessed by monitoring the release rate of molecules encapsulated in bulk hydrogels. We conducted diffusion experiments using the same setup for all investigated molecules, including inflammatory markers and model molecules. Similar release experiments were performed with ionically cross-linked alginate hydrogels, which have previously been demonstrated to maintain cell bioactivity both in vitro and in vivo [21]. After 7 days, 74% of the fluorescein encapsulated in alginate-based SPAAC hydrogels was released, and 49% of fluorescent BSA was released from the same hydrogel. We also noted a burst release within the first 6 hours for fluorescein, followed by a plateau. In contrast, fluorescent BSA continued to be released for up to 2 days before reaching a plateau (Fig 2b). The results indicated similar diffusion profiles, with a burst release over the first 6 hours and a higher fluorescein release rate of 57 %, compared to the 19 % fluorescent BSA released after 7 days in the ionic alginate hydrogel. When comparing the two hydrogels, it is noteworthy that the release rate of model molecules is higher in the alginate-based SPAAC hydrogel compared to the ionic alginate control. We then studied the release of two pro-inflammatory cytokines from alginate-based SPAAC hydrogels (Fig 2c). Following a study period of 3 days, it was found that only 9 % of IFN-γ was released, whereas the TNF-α release rate was higher, reaching 46 %. Regarding the ionically cross-linked alginate hydrogel, the two cytokines showed similar release rates, with IFN-γ and TNF-α reaching release plateau of 41 % and 42 %, respectively. The difference in IFN-γ release in alginate-based SPAAC hydrogels could be attributed to potential interactions between the network and the molecule and possible differences in their charges.

**Figure 2:**
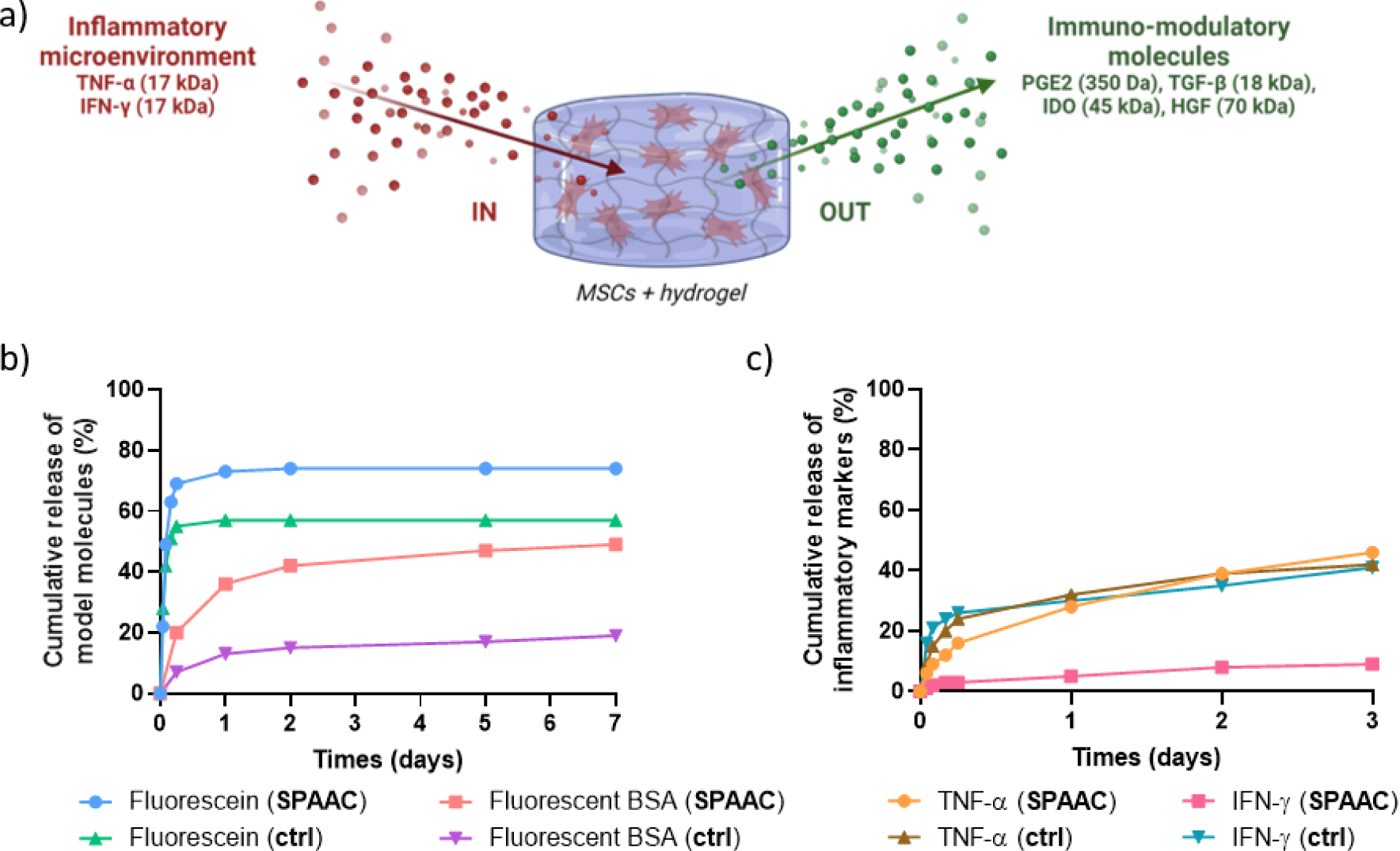
Diffusion properties of alginate-based SPAAC hydrogels and secretory activity of microencapsulated hASCs. a) Schematic representation illustrating the diffusion of molecules of interest through a hydrogel in an inflammatory microenvironment. Cytokines, such as TNF-α and IFN-γ, enter the hydrogel to trigger the activation of encapsulated MSCs, and in response this stimulus, encapsulated MSCs secrete immunomodulatory molecules, such as PGE2, TGF-β, IDO and HGF, that exit the hydrogel and exert their effect on surrounding cells; b) Diffusion profiles of model molecules, fluorescein and fluorescent BSA, that mimic the molecular weight of secretory markers (N=1 or 2, n=4) in PBS (ctrl = ionic alginate); c) Diffusion profiles of inflammatory markers, TNF-α and IFN-γ (N=1 or 2, n=4) in PBS (ctrl = ionic alginate).

### 3.4 Micro-encapsulation of hASCs in alginate-based SPAAC hydrogels

To increase the exchange surface between the inside and outside of the hydrogels, we prepared alginate-based SPAAC microgels using a micromolding technique (Fig. 3a). Cylindrical PDMS micromolds of 150 µm in diameter and 100 µm in height were used. We characterized the shape and size of the incubated microgels over time using light microscopy (Fig. 3b). The hydrogels were incubated in complete cell culture medium at 37°C for 21 days. Our results showed that alginate-based SPAAC microgels had a mean diameter of 168 ± 7 µm at D0, 165 ± 6 µm at D1, 166 ± 7 µm at D3, 168 ± 5 µm at D7, 165 ± 15 µm at D14 and 166 ± 15 µm at D21, with no statistical difference over time (Fig. 3c). This finding indicates that the alginate-based SPAAC microgels remained stable throughout the observation period. Next, human adipose-derived mesenchymal stromal cells (hASCs) were encapsulated in microgels using the previously outlined procedure. The alginate-based SPAAC hydrogel was first loaded with cells using a cell density of 3 million cells per mL of pre-hydrogel solution (1% w/v, BCN/N_3_ molar ratio 1:1.4). Following a centrifugation step to enhance cell encapsulation in the micromolds and a 1-hour cross-linking process at 37°C, the hASCs-loaded microgels were harvested and observed under light microscopy (Fig. 3d). Immediately after the procedure, we observed that the hASCs were located within the microgels and homogeneously distributed. We repeated the microgel stability assessment on cell-loaded microgels by placing them in complete cell culture medium at 37°C for 14 days before light microscopy analysis. Their mean diameter was 146 ± 7 µm at D0, 141 ± 10 µm at D1, 139 ± 8 µm at D3, 142 ± 9 µm at D7, and 142 ± 12 µm at D14 (Fig. 3e). The cell-loaded microgels maintained their diameter and shape, with no significant changes in the mean diameter in 14 days.

**Figure 3:**
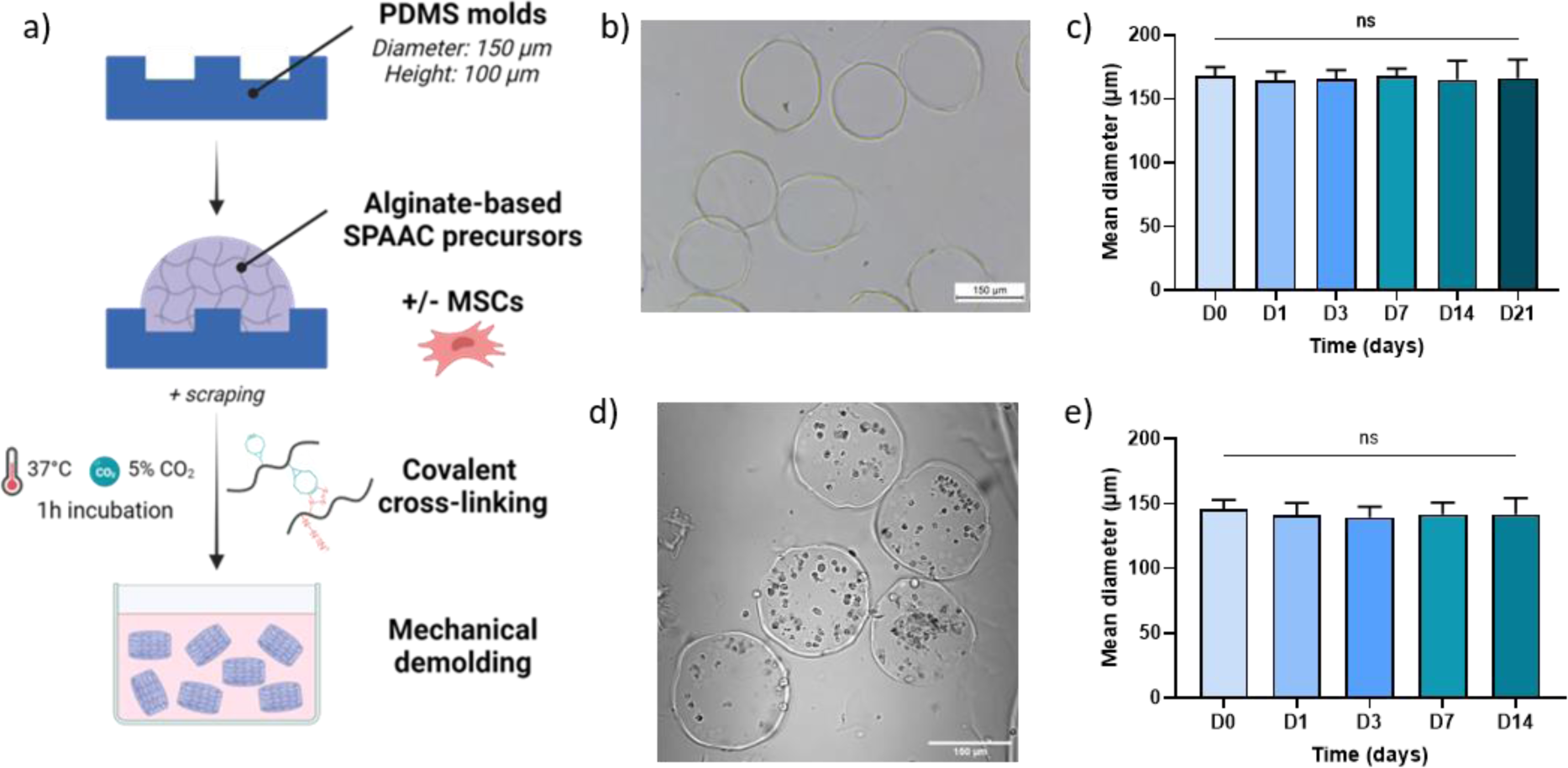
Cell encapsulation in alginate-based SPAAC microgels. a) Schematic representation illustrating the micromolding procedure +/- MSCs; b) Microgels observed under light microscopy, scale bar 150 µm; c) Stability of microgels over 21 days (N=3, n=100); d) Cell-loaded microgels observed under light microscopy, scale bar 150 µm; e) Stability of cell-loaded microgels over 14 days (N=1, n=25).

### 3.5 Cytocompatibility of alginate-based SPAAC hydrogels on hASCs

The suitability of alginate-based SPAAC microgels for 3D cell culture was then examined. The viability of encapsulated hASCs was assessed both immediately after microencapsulation and throughout a 14-day culture period, demonstrating that more than 90% of the cells remained viable within the microgels (Fig. 4a and 4d). Furthermore, using the CCK-8 and PicoGreen assays, we observed a stable metabolic activity of the microencapsulated hASCs (Fig. 4b), along with a constant DNA concentration of 109 ± 10 ng/mL (Fig. 4c), highlighting the absence of hASCs proliferation in 3D. In contrast, when examining cells cultured in 2D, we observed an increase in metabolic activity and DNA concentration, from 230 ± 22 ng/mL at D0 to 1068 ± 113 ng/mL at D14, indicating cell proliferation during the 14-day culture period. To observe the cells and their behavior in the alginate-based SPAAC microgels, Live/Dead with TO-PRO- 3 staining (Fig. 4d) and Alexa Fluor 568 Phalloidin with TO-PRO-3 staining (Fig. 4e) were performed over the 14 days of culture. The resulting data were acquired using confocal microscopy. We observed changes in cell behavior over the culture period, characterized by modifications in the cell cytoskeleton. Further investigation, including electronic microscopy, will be necessary to understand this phenomenon better.

**Figure 4:**
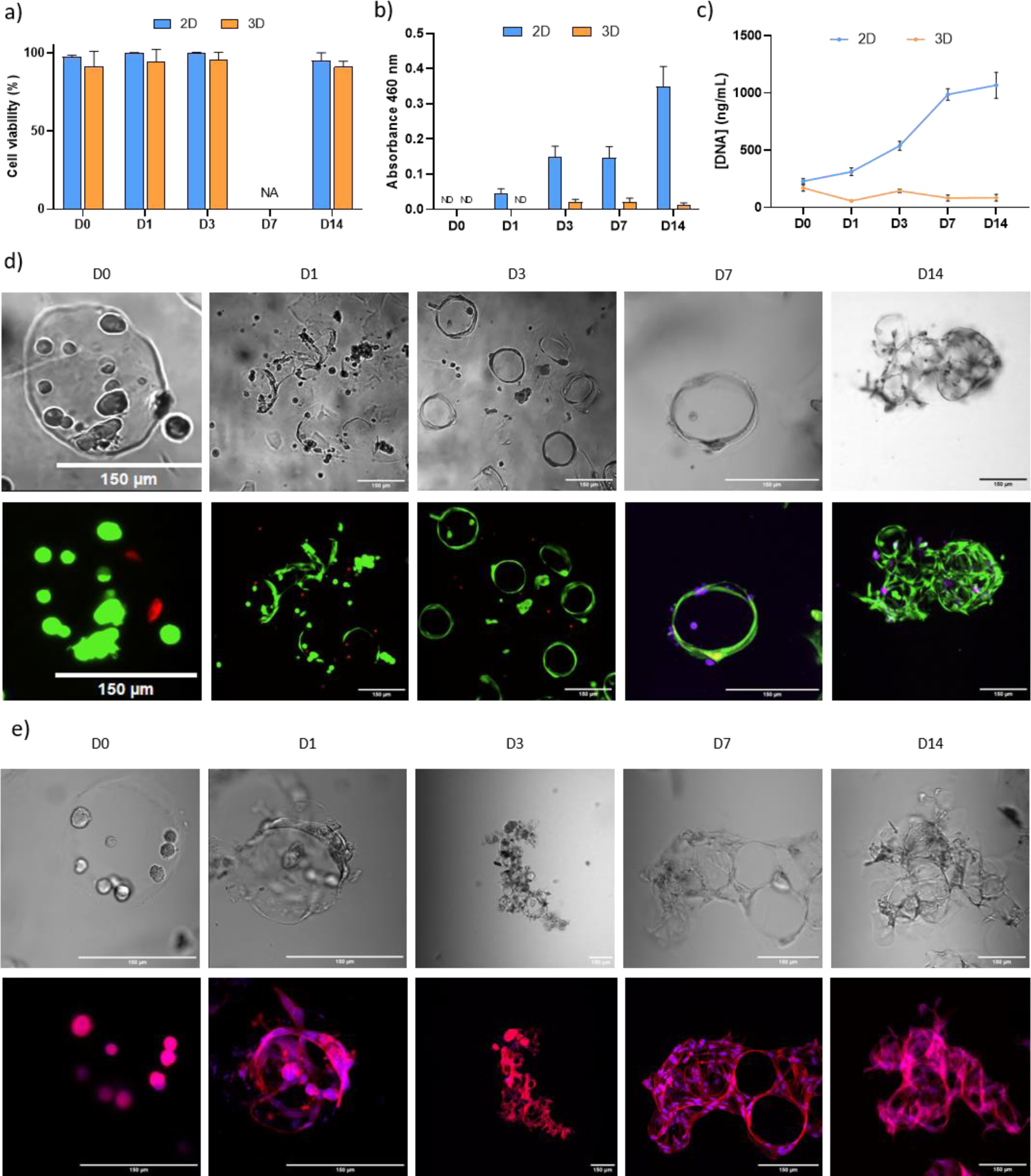
Cytocompatibility of alginate-based SPAAC microgels on hASCs. a) Cell viability, b) metabolic activity (absorbance 460 nm) and c) DNA quantification of hASCs in 2D or encapsulated in alginate-based SPAAC microgels, during 14 days of cell culture (N=1, n=3); d) LIVE/DEAD with TO-PRO-3 staining images and e) Alexa Fluor 568 Phalloidin with TO-PRO-3 staining images of hASCs microencapsulated, during 14 days of cell culture, scale bar 150 µm (N=1, n=3).

### 3.6 Biological activity of microencapsulated hASCs

#### Secretory activity

After demonstrating that the cells were viable in alginate-based SPAAC microgels and that some molecules of interest diffused through these hydrogels, we investigated the immunomodulatory properties of microencapsulated hASCs. We then evaluated the secretory properties of microencapsulated hASCs. Cells were treated with pro-inflammatory factors, i.e., TNF-α and IFN-γ, each at a concentration of 20 ng/mL, for 72 hours as previously described [21], [29], before assessing the secreted IDO activity and the concentrations of secreted PGE2, HGF, and TGF-β in the supernatants (Fig. 5a). The results were normalized to the total cell number in each sample. The treatment with TNF-α and IFN-γ induced a significant increase in the IDO activity compared to the control condition, i.e., unstimulated microencapsulated hASCs, increasing from 10.3 ± 6.7 pM to 14.6 ± 4.7 pM (Fig 5b). In addition, PGE2 concentration was also significantly increased in the microencapsulated hASCs after 72 hours of TNF-α and IFN-γ treatment compared to the control, increasing from 6 ± 3 fg/mL to 410 ± 208 fg/mL (Fig 5c). The concentration of HGF was significantly reduced when hASCs were treated with TNF-α and IFN-γ compared to the control, with concentrations decreasing from 28 ± 13 fg/mL to 21 ± 11 fg/mL (Fig 5c). This decrease in HGF concentration could be partly attributed to the high MW of this factor, i.e., 70 kDa. The concentration of TGF-β was not detectable in this experiment.

**Figure 5:**
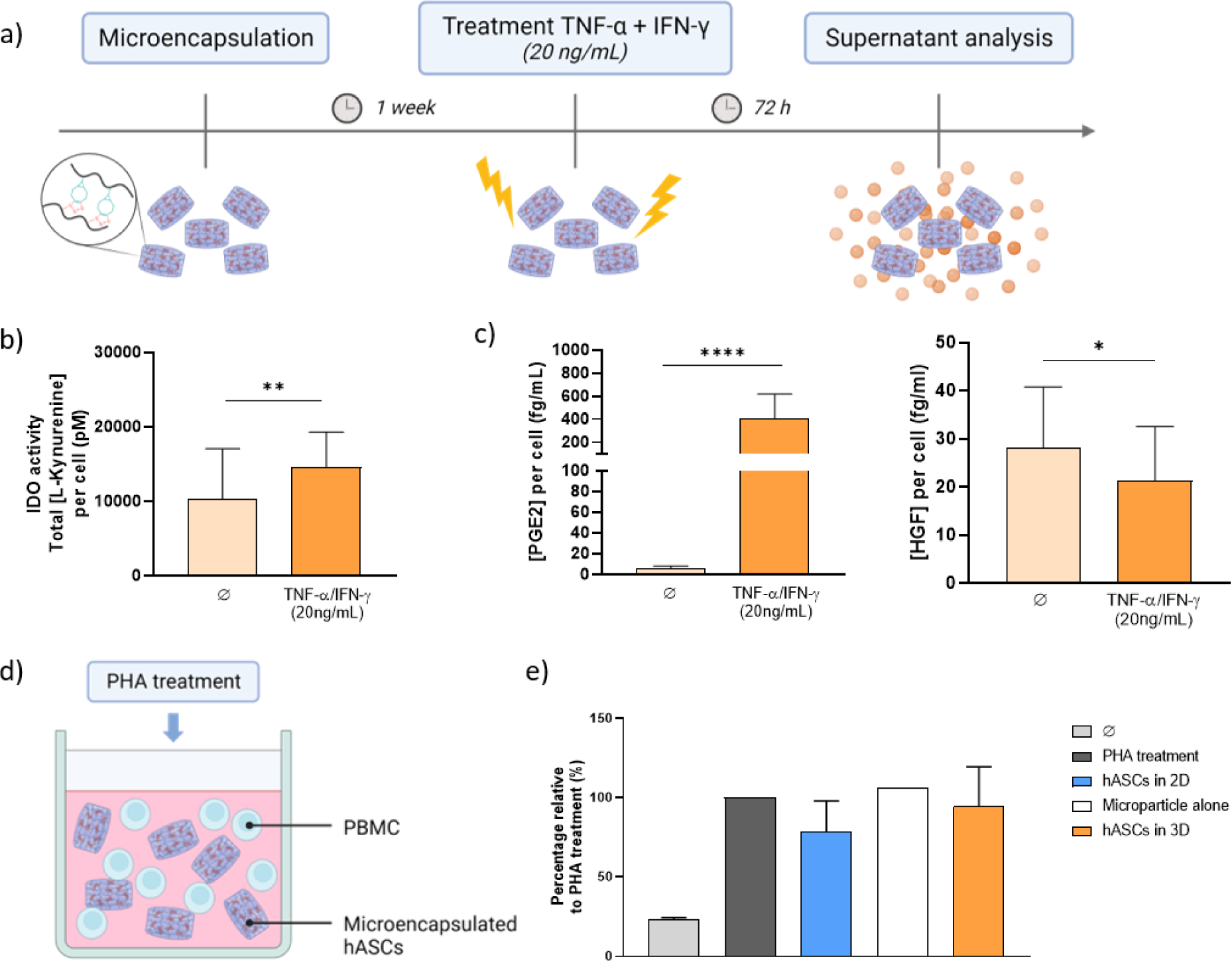
Secretory and immunomodulatory activities of microencapsulated hASCs. a) Schematic representation illustrating the treatment protocol of hASCs microencapsulated; In vitro secretion of hASCs of b) IDO enzyme and c) PGE2 and HGF, dosed in supernatants and normalized to the total cell number, after 72 hours of TNF-α/IFN-γ treatment (N=3 human donors, repeated 2 times on each donor, n=5; * p<0.05, ** p<0.01, **** p<0.0001, unpaired t-test); d) Schematic illustrating the co-culture protocol; e) In vitro immunomodulation of microencapsulated hASCs on lymphocytes (N=3 human donors; Kruskal- Wallis test).

#### Immunomodulatory activity

To address the effect of encapsulation on the immunosuppressive properties of ASCs, we cultured PHA-activated PBMCs in the presence of hASCs, cultured in 2D or microencapsulated, at a PBMCs / hASCs ratio of 10:1. PBMC from 3 donors were pooled and added into a 96-wells plate with MLR medium. hASCs from 3 donors were co-cultured for 4 days (Fig 5d). When hASCs were cultured in 2D, we observed a trend toward an inhibitory effect on T cell proliferation (Fig 5e). This tendency was confirmed when hASCs were encapsulated. However, in both cases, differences were not significant. Moreover, when microgels alone were added into the MLR medium, the proliferation of CD3+ cells was not modified.

## 4 Discussion

Alginate-based SPAAC hydrogels are obtained in a one-step synthesis through the covalent cross-linking of two initial precursors, alginate-BCN and alginate-N_3_. This cross-linking process respects the principles of click and bioorthogonal chemistry, i.e., physiological medium and temperature, no additional reagents, no by-products, and high yields. The molecular weight of alginate, measured at ≈ 67 kDa, enabled filter sterilization of the precursors and, therefore, easy to use for 3D cell culture applications. In the scientific literature, several systems have been developed for the formation of alginate hydrogels by covalent cross-linking, including the thiol-ene reaction [32] or the inverse electron-demand Diels-Alder (IEDDA) reaction [33], [34]. However, these reactions have certain drawbacks that limit their use in the biological and medical fields. The thiol-ene reaction can lead to the formation of unstable products. In contrast, IEDDA involves the production of nitrogen gas as a by-product, leading to the formation of bubbles in hydrogels. For these reasons, the SPAAC click reaction was developed to fulfill the criteria of both click chemistry and bioorthogonal chemistry. To the best of our knowledge, only one paper in the literature describes the covalent cross-linking of alginate via the SPAAC reaction. In this study, alginate is functionalized with azide groups and cross-linked in the presence of dendrimers carrying tetrabicyclononynes groups to form covalent hydrogels [30]. However, using a complex synthetic polymer, i.e., the PAMAM dendrimer, limits its use for a medical application [30]. The research most closely related to our system was conducted in our own laboratory, where a different polysaccharide, hyaluronic acid, was functionalized with a BCN or an N_3_. Therefore, our study appears as the first investigation to document the functionalization of alginate with BCN or N_3_ molecules. The stability of alginate-based SPAAC hydrogels was investigated in both PBS and DMEM culture medium. These investigations indicated that the hydrogels remained stable for 2 months in culture conditions as long as the hydrogel was cross-linked for at least 1 hour at 37°C. It would be interesting to explore the stability of these hydrogels in the synovial fluids of OA patients, which mimic a more realistic OA environment, while considering that their composition is highly variable.

In this study and for developing an OA cell therapy, it was crucial to generate micro-sized hydrogels to enable the intra-articular injection of alginate-based SPAAC hydrogels through a needle. We demonstrated the feasibility of producing alginate-based SPAAC microgels using the micromolding technique developed in our laboratory and previously used to micromold ionically cross-linked alginate hydrogels [21]. We have successfully generated microgels of cylindrical shape with reproducible size (average diameter of 166 µm) that were easily removed from the micromolds. However, the cross-linking time to form stable alginate-based SPAAC microgels, 1 hour at 37°C, is longer than the cross-linking time to create ionic alginate microgels, which is 5 minutes at room temperature. Therefore, in the future, it would be interesting to explore potential modifications of the alginate-based SPAAC hydrogel formulation to reduce the cross-linking time, by increasing the alginate concentration or molecular weight.

For cell therapy applications, hASCs were successfully encapsulated in alginate-based SPAAC microgels. The cytocompatibility of the microgels was confirmed both immediately after microencapsulation and throughout a 14-day culture period, with more than 90 % viable cells within the microgels, along with constant metabolic activity and DNA concentration. We observed an interesting phenomenon that led to an unexpected morphology of the microencapsulated cells. Surprisingly, the cells initially present in the microgels on the day of encapsulation were no longer found at the center of the microgels after 3, 7, and 14 days of culture but rather at their periphery. Various hypotheses have been proposed to explain this phenomenon, including the possibility of cell mobility within the microgels, potential interactions between the cells and the functional groups present on the polymer chains, or the adherence of non-encapsulated cells to the microgel surface. The latter hypothesis was refuted by running additional experiments where a cell suspension was incubated with empty microgels for up to 7 days. We confirmed that cells could not adhere to the surface of the microgels (data not shown). Given the absence of existing literature references regarding this specific morphology in the context of alginate-based SPAAC hydrogels, we can only make assumptions. Conducting a “time-lapse” analysis of the samples could provide answers to these questions.

Molecular diffusion is vital to a successful hydrogel-assisted immunomodulatory cell therapy. The cytokines produced in an inflammatory microenvironment must easily reach the hASCs in the microgels. Then, the secreted molecules must be able to diffuse through the hydrogel toward the culture medium (in vitro) or the local microenvironment (in vivo) and finally reach their target cells. We chose to perform diffusion experiments using a similar setup for all the molecules investigated, including both inflammatory and secretory markers. To investigate the diffusion properties of alginate-based SPAAC hydrogels, molecules of various molecular weights, charges, and hydrophilic properties were selected, including fluorescein, fluorescent BSA, TNF-α, and IFN-γ. Our experiments revealed that the molecular weight of the molecule and the nature of the hydrogel network influence the diffusion of model molecules and pro-inflammatory cytokines. We chose two fluorescent model molecules covering a wide range of molecular weights to carry out preliminary experiments on the diffusion properties of alginate-based SPAAC hydrogels: a low-MW molecule (fluorescein, 370 Da) and a high-MW protein (fluorescent BSA, 66 kDa). We observed that both model molecules successfully diffused through our alginate-based SPAAC hydrogels. These results provide insights regarding the effect of the molecular weight on molecular diffusion, which may then allow to partially anticipate the expected diffusion profiles of molecules of interest, including PGE2 (350 Da), the IDO enzyme (45 kDa), HGF (70 kDa) and TGF-β (18 kDa). Next, we investigated the release of two pro-inflammatory cytokines through our alginate-based SPAAC hydrogels, TNF-α and IFN-γ, both having identical MW (17 kDa). The difference in IFN-γ release could be attributed to possible interactions between the hydrogel network and the charged molecule. In future experiments, one possible approach could be to reduce the polymer concentration to increase the network mesh size, thereby promoting the diffusion of high-MW molecules through the alginate-based SPAAC hydrogel.

Following the confirmation of the diffusion of some molecules of interest through these hydrogels, we investigated the ability of microencapsulated hASCs to secrete immunomodulatory factors in response to an inflammatory environment. The cells were exposed for 72 hours to pro-inflammatory factors, i.e., TNF-α and IFN-γ, as previously reported [21], [29]. The IDO activity and the concentration of PGE2 were significantly higher when the microencapsulated hASCs were treated with pro-inflammatory factors compared to the untreated microencapsulated cells. These results demonstrate that microencapsulated MSCs remain sensitive to pro-inflammatory stimuli and maintain their anti-inflammatory properties. It is important to note that we observed a weaker treatment effect on IDO activity, which may be related to the limited IFN-γ diffusion capacity. Indeed, the association between IFN-γ and the expression of the IDO gene has been demonstrated in various human cells, notably in synovial cells. When IFN-γ binds to its receptor, it triggers the activation of the gene responsible for producing IDO [35]. In the case of alginate-based SPAAC hydrogels, IFN-γ stimulation is probably insufficient due to limited diffusion. Moreover, the concentration of HGF was significantly reduced when hASCs were treated with TNF-α and IFN-γ, compared to the untreated microencapsulated cells. This decrease in HGF concentration could be partly attributed to its high MW of 70 kDa.

We established direct co-culture experiments to assess the immunomodulatory effects of microencapsulated hASCs on specific target cells, such as lymphocytes. Following 4 days of co-culture, we quantified the proliferation of CD3+ T cells by measuring the corresponding decrease in cell fluorescence through flow cytometry analysis. The findings demonstrate a trend toward immunosuppression of CD3+ cells in the presence of hASCs, whether grown in 2D or microencapsulated. One of the drawbacks of this study is the high variability of hASCs, which can be quite heterogeneous in terms of phenotype and differentiation potential. To address this intrinsic cellular variability, one potential approach to consider would be to standardize hASCs.

## 5 Conclusions

The present study describes the generation of alginate-based SPAAC micro-sized hydrogels using a combination of alginate-BCN and alginate-N_3_. As far as we know, the micromolding of alginate-based SPAAC hydrogels has not been attempted before. Micromolded alginate-based SPAAC hydrogels were stable in vitro and cytocompatible. Microencapsulated hASCs maintained their secretory and immunomodulatory properties, confirming the potential of the alginate-based SPAAC microgels for cell therapy

## Acknowledgments

The authors thank Dr. F. Lejeune (Clinique Breteche, Nantes) for harvesting the human lipoaspirates. The authors acknowledge the SC3M platform from the Inserm/NU/ONIRIS UMR1229 RMeS Laboratory (SFR Bonamy, BioCore, Inserm UMS 016, CNRS UAR 3556, Nantes, France), the IBISA MicroPICell facility (Biogenouest), member of the national infrastructure France-Bioimaging supported by the French national research agency (ANR-10- INBS-04), and the Cytocell - Flow Cytometry and FACS core facility (SFR Bonamy, BioCore, Inserm UMS 016, CNRS UAR 3556, Nantes, France), member of the Scientific Interest Group (GIS) Biogenouest and the Labex IGO program supported by the French National Research Agency (n°ANR-11-LABX-0016-01).

This study was supported by grants from the Fondation de l’Avenir pour la Recherche Médicale Appliquée (AP-RM-18-005), the RFI Bioregate program of Pays de la Loire Region within the framework of the BOROHYDROGEL project, the Nantes Excellence Trajectory (NExT) funding scheme within the framework of the SHELBY project and the METAB-OA project (ANR-20-CE18-0014).

## Supplementary data

**Suppl. Figure S1:**
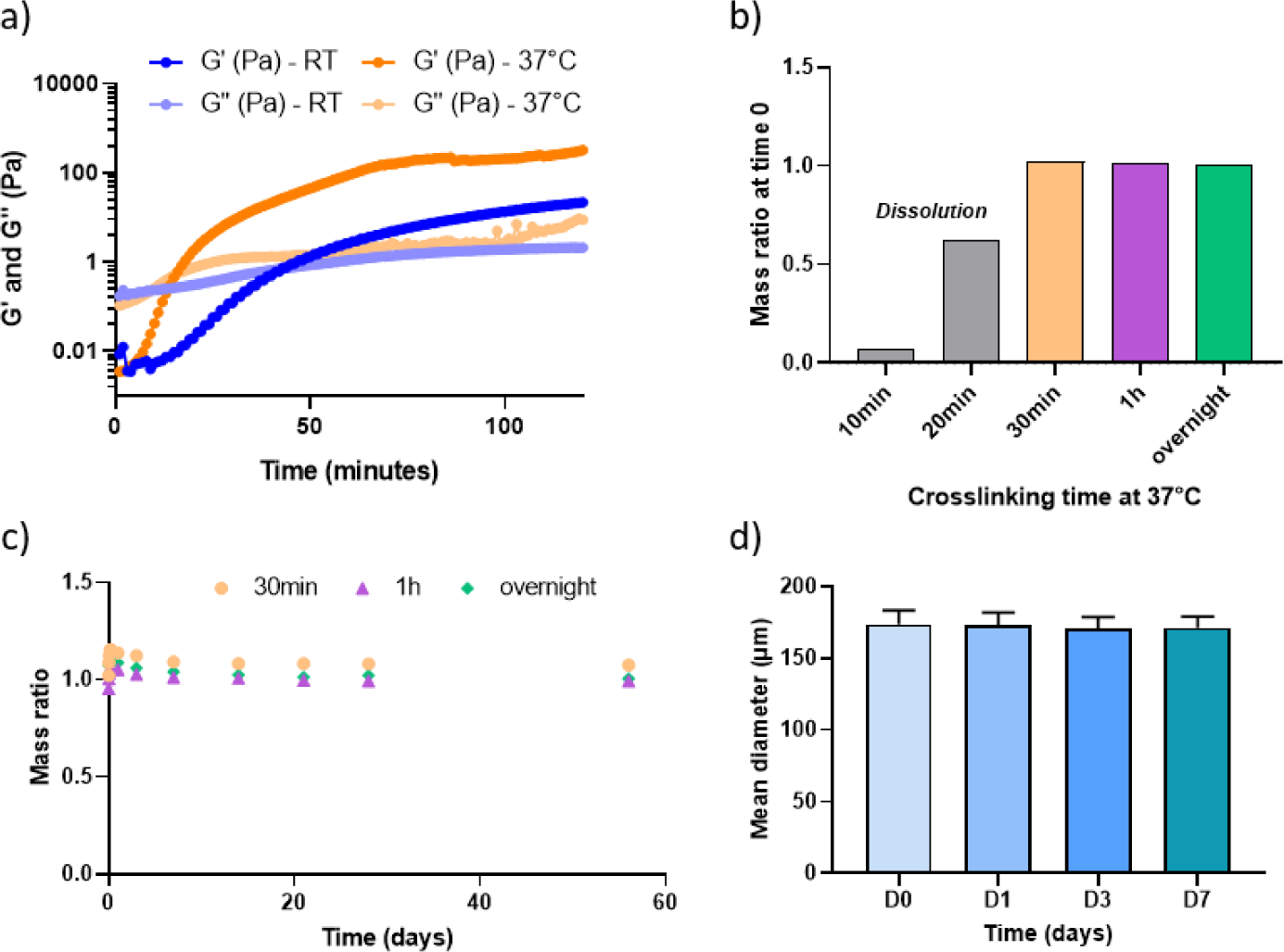
Physicochemical characterization of alginate-based SPAAC hydrogels in PBS. a) Time sweep experiments representing the evolution of G’ and G’’ as a function of time and temperature in PBS (N=3, n=1); The influence of cross-linking time on hydrogel mass ratio on stability : b) at short-term (N=1, n=1) and c) over 2 months (N=1, n=3) in PBS; d) Stability of microgels over 21 days (N=1, n=100 ).

**Suppl. Figure S2:**
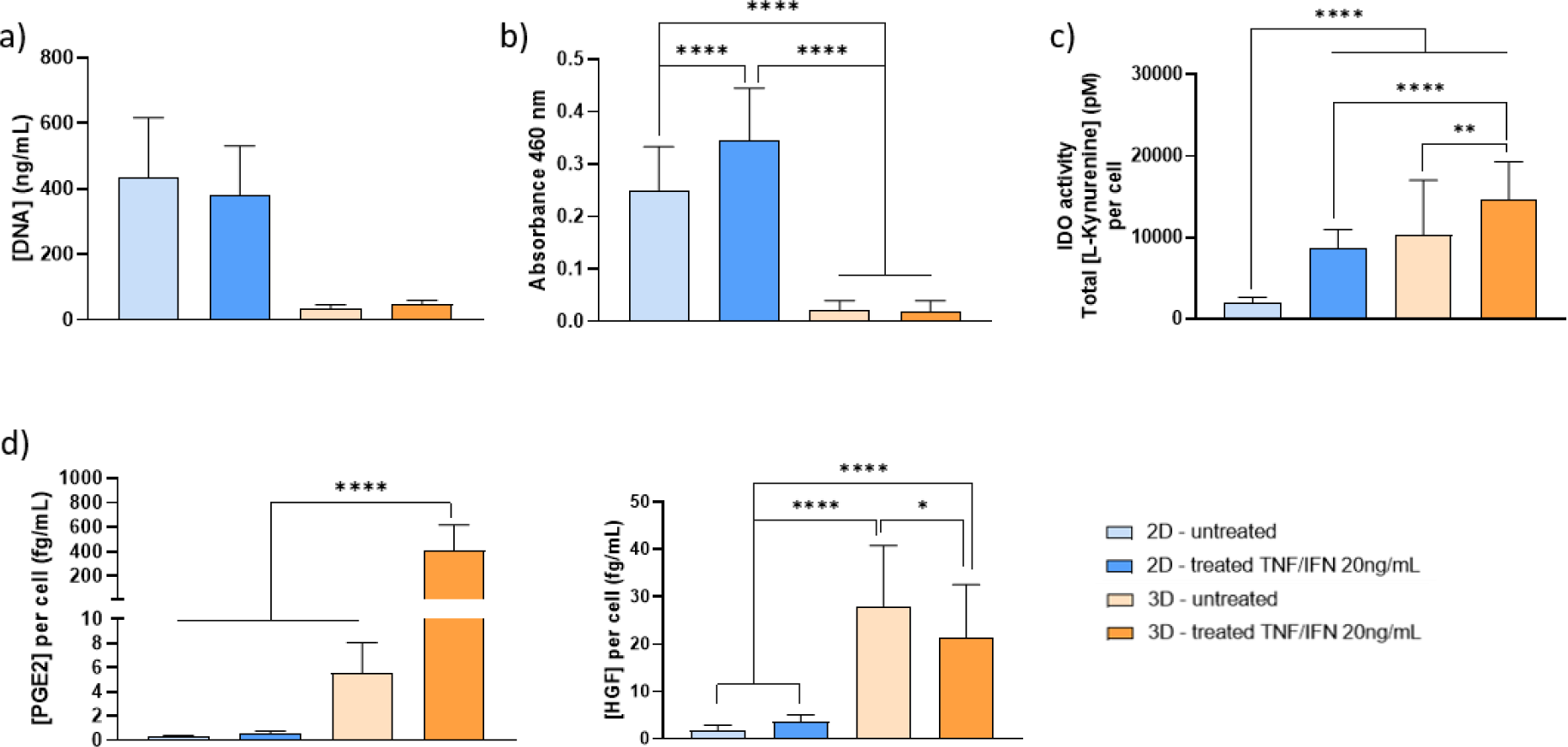
Secretory activity of microencapsulated hASCs. a) DNA quantification, b) metabolic activity (absorbance 460 nm) and in vitro secretion of hASCs in supernatants of c) IDO enzyme and d) PGE2 and HGF, normalized to the total cell number, after 72 hours of TNF-α/IFN-γ treatment (N=3 human donors, n=2; ** p<0.01, *** p<0,001, *** p<0.0001, ANOVA).

**Suppl. Table 1:**
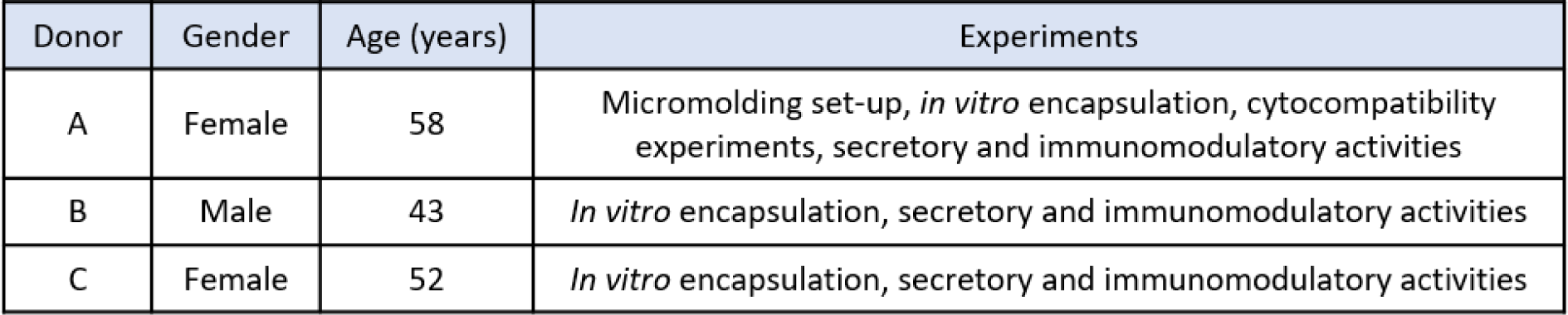
Three different donors of human adipose-derived mesenchymal stromal cells (hASCs).

## Notes

### Competing Interest Statement

The authors have declared no competing interest.

### Summary of Updates

Update information regarding the confocal imaging of samples

