## Supplementary for "Microencapsulation of mesenchymal stromal cells in covalent alginate hydrogels for cell therapy"

### Supplementary data

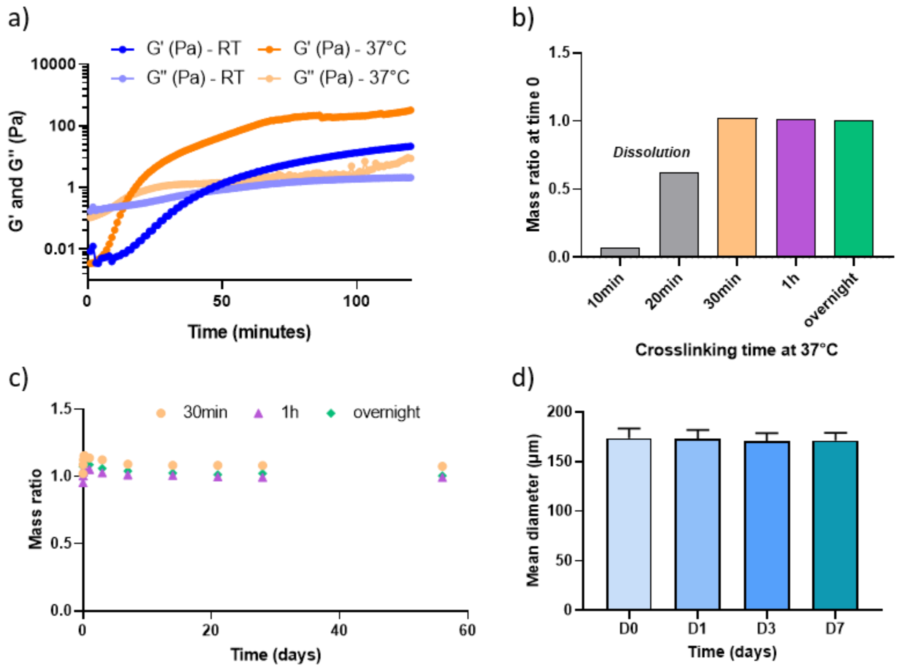


*<u>Suppl. Figure S1: Physicochemical characterization of alginate-based SPAAC hydrogels in PBS.</u> a) Time sweep experiments representing the evolution of G' and G'' as a function of time and temperature in PBS (N=3, n=1); The influence of cross-linking time on hydrogel mass ratio on stability : b) at short-term (N=1, n=1) and c) over 2 months (N=1, n=3) in PBS; d) Stability of microgels over 21 days (N=1, n=100 ).*

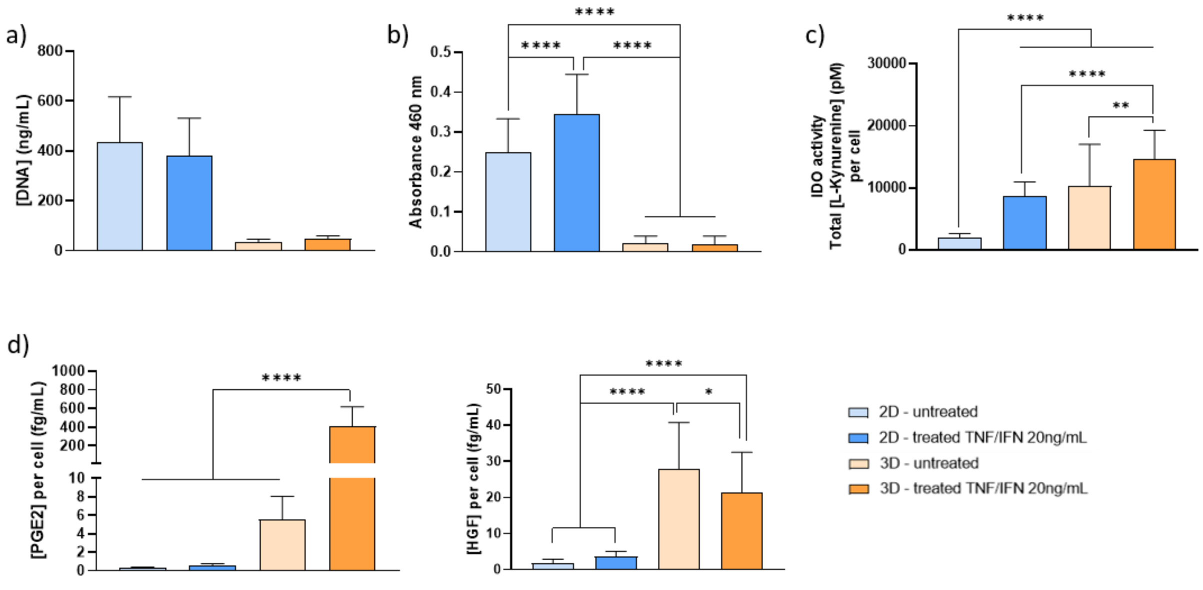


*<u>Suppl. Figure S2: Secretory activity of microencapsulated hASCs.</u> a) DNA quantification, b) metabolic activity (absorbance 460 nm) and in vitro secretion of hASCs in supernatants of c) IDO enzyme and d) PGE2 and HGF, normalized to the total cell number, after 72 hours of TNF-α/IFN-γ treatment (N=3 human donors, n=2; ** p<0.01, *** p<0,001, *** p<0.0001, ANOVA).*

| Donor | Gender | Age (years) | Experiments |
|---|---|---|---|
| A | Female | 58 | Micromolding set-up, *in vitro* encapsulation, cytocompatibility experiments, secretory and immunomodulatory activities |
| B | Male | 43 | *In vitro* encapsulation, secretory and immunomodulatory activities |
| C | Female | 52 | *In vitro* encapsulation, secretory and immunomodulatory activities |

*Suppl. Table 1: Three different donors of human adipose-derived mesenchymal stromal cells (hASCs).*
